# Network-based analysis of glioblastoma identifies patient communities and cluster-specific biomarkers

**DOI:** 10.64898/2026.04.30.721983

**Authors:** Nicoleta Siminea, Daniela Florea, Mihaela Păun, Andrei Păun, Ion Petre

## Abstract

Glioblastoma is an aggressive and highly heterogeneous brain tumor with poor prognosis despite multimodal treatment strategies. Understanding the molecular diversity of the disease is essential for improving tumor stratification and identifying potential therapeutic targets. In this study, we investigate whether network-based analysis can reveal biologically meaningful subgroups of glioblastoma tumors. Using RNA sequencing and mutation data from the TCGA-GBM cohort, we constructed patient-specific protein–protein interaction networks based on genes that are differentially expressed or harbor somatic mutations. These networks capture the molecular alterations associated with individual tumors within the context of the human interactome. We then derived similarities between tumors using a binary representation of network nodes and the Jaccard similarity metric, enabling the construction of a patient similarity graph. Community detection algorithms (Louvain and Leiden) were applied to this graph to identify clusters of tumors with similar molecular network profiles. Our analysis revealed six tumor communities characterized by distinct gene compositions and enriched biological processes. For each community, we identified candidate biomarkers and network hubs that may represent potential therapeutic targets. Several of the identified genes correspond to known drug targets, while others represent potential candidates for further investigation.

These results illustrate how integrating molecular alterations with network-based modeling can help stratify glioblastoma tumors and uncover molecular mechanisms that may guide the development of more personalized therapeutic strategies.

## 1 Introduction

Glioblastoma is the most aggressive primary brain tumor in adults and is associated with a median overall survival of approximately 15 months [1] despite multimodal treatment strategies that typically include surgical resection, radiotherapy, and chemotherapy. The poor prognosis is partly due to the high molecular and clinical heterogeneity of the disease, which complicates both diagnosis and treatment selection [2]. As a result, identifying molecular patterns that distinguish subgroups of patients remains an important challenge in glioblastoma research. Improved stratification of tumors could facilitate the identification of biomarkers and support the development of more targeted therapeutic strategies.

Our objective was to identify patterns within glioblastoma in order to better understand the molecular mechanisms that contribute to its development and progression. Ultimately, we aim to improve tumor stratification and support the identification of more appropriate therapeutic strategies for patients affected by this disease. A substantial body of research has investigated the molecular heterogeneity of glioblastoma. A landmark study by [3] applied hierarchical clustering to genomic data and identified four biologically relevant glioblastoma subtypes.

In this work, we adopt a different perspective by applying community detection methods to identify groups of tumors with similar molecular characteristics. In particular, we construct a similarity network of tumor samples and detect communities within this network. This approach differs from studies such as [4], where communities were identified from gene co-expression networks across several cancer types, including glioblastoma. Co-expression network approaches have also been widely used for biomarker discovery, especially through Weighted Gene Co-expression Network Analysis (WGCNA), as illustrated in studies such as [5] and [6].

Other studies have explored glioblastoma heterogeneity using alternative data modalities. For example, [7] identified two glioblastoma clusters derived from diffusion and perfusion MRI similarities, highlighting the potential of imaging-based clustering for capturing intratumoral heterogeneity and improving prognostic assessment. At the cellular level, [8] used single-cell RNA sequencing data and hierarchical clustering to identify five malignant cell clusters and their corresponding prognostic modules.

In this study, we conduct an *in silico* analysis that integrates differentially expressed and mutated genes to construct protein–protein interaction (PPI) networks for individual tumor samples. These networks are characterized using several network centrality measures. From the resulting networks, we derive a sample-similarity network and apply community detection algorithms to identify six distinct modules. Each module is further characterized by its specific gene composition, from which we select candidate genes with high module specificity for potential biomarker evaluation. For each module, we identify the three most significant biological processes uniquely associated with that module through enrichment analysis. These processes reflect distinct biological mechanisms that may contribute to glioblastoma progression, and the associated genes are mapped to our candidate lists.

Finally, for each module we construct a consensus network that integrates the interactions observed across samples belonging to that module. These networks are analyzed using centrality metrics to identify key hub genes. Notably, several hubs overlap with our candidate biomarkers and are involved in critical biological processes. Some of these genes are targets of existing drugs, while others represent potential candidates for future therapeutic investigation.

Our approach combines molecular alterations with network-based analysis to identify groups of tumors with similar molecular characteristics. First, we construct patient-specific protein–protein interaction networks using genes that are either differentially expressed or harbor somatic mutations. These networks are then converted into binary gene-presence representations, allowing similarities between tumors to be quantified using the Jaccard metric. Based on these similarities, we construct a patient similarity graph and apply community detection algorithms to identify clusters of tumors. For each detected community, we characterize its gene composition, perform functional enrichment analysis, and identify candidate biomarkers and central network hubs. This framework enables the identification of glioblastoma subgroups together with potential molecular targets associated with each group. Our results show that network-based stratification can reveal biologically meaningful tumor communities and provide candidate biomarkers and therapeutic targets associated with each subgroup. Our approach differs from existing network-based analyses in that it derives tumor communities from a patient similarity network constructed from patient-specific protein–protein interaction networks. This strategy allows molecular alterations to be interpreted in their network context while simultaneously capturing similarities between tumors at the systems level.

Our contributions are threefold: (i) we introduce a network-based framework for stratifying glioblastoma tumors using patient-specific protein–protein interaction networks; (ii) we construct a tumor similarity graph and identify communities using modularity-based community detection; and (iii) we characterize the resulting communities through functional enrichment, biomarker identification, and network hub analysis, revealing potential therapeutic targets.

## 2 Methods

Our study is a computational analysis that builds upon well-established, publicly available datasets. It applies network-based approaches, including the use of network centrality measures and community detection algorithms, combined with enrichment analysis to explore the underlying biological mechanisms.

### 2.1 Datasets

The datasets used in our study comprise genetic data taken from individual cases as well as protein-protein interaction data.

The data for individual cases were obtained from the GDC database, specifically from the TCGA-GBM project [9]. Our study was made on 134 tumor cases and 5 healthy cases. For each tumor case, RNA sequencing (RNA-seq) results and mutation data, when available, were included in the analysis. The retrieved RNA-seq data were filtered to exclude samples derived from recurrent tumors and those with annotations that could potentially affect the genetic data (e.g., prior treatment or a different type of cancer). For each case, we retrieved somatic mutations from the same database that were predicted to affect protein function or, in some cases, protein expression. Using this criterion, mutations with a high predicted impact according to the Variant Effect Predictor (VEP) were identified in 127 of the tumor samples.

The protein-protein interaction dataset was taken from [10] and includes the human interactome.It comprises more than 320 000 interactions between human proteins.

### 2.2 Differential expression analysis

Differential expression analysis was performed by comparing the cancer group to the control group using the glmQLFTest function from the edgeR package [11]. Results were filtered using a false discovery rate (FDR) threshold of 0.05 and an absolute log2 fold change (|log2FC|) greater than 1. Next, we identified up-regulated genes at the individual level by selecting those with logCPM values above the sample mean and that were also up-regulated at the cohort level. Similarly, down-regulated genes were defined as those under-expressed relative to the sample mean and down-regulated at the cohort level. We converted them from Ensembl names to Hugo names using BioMart [12].

### 2.3 Networks construction and analysis

The subsequent steps of the analysis are presented in the Figure 1.

**Figure 1:**
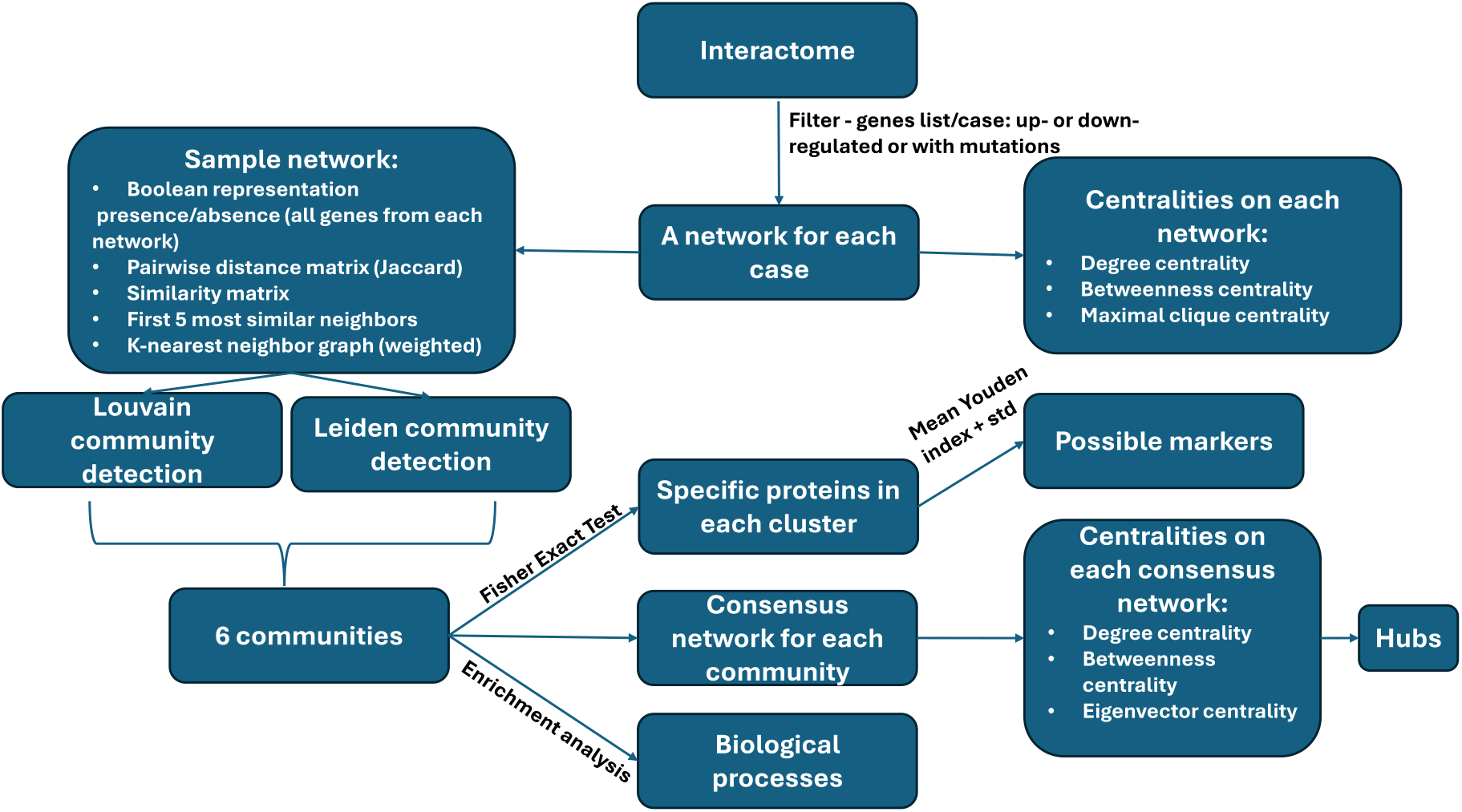
The applied methodology.

For each case, we constructed an undirected protein–protein interaction (PPI) network using NetworkX [13]. From the human interactome, we extracted all interactions in which both the interacting proteins were present in the case-specific gene list. This approach allowed us to define a a network for each case, comprising proteins encoded by differentially expressed genes as well as genes harboring mutations.

In biological networks, some of the most relevant nodes are typically those with high degree, high betweenness centrality, or high maximal clique centrality. High-degree nodes interact with many other nodes, while nodes with high betweenness centrality often act as bridges connecting different parts of the network. Maximal clique centrality identifies nodes located in densely connected regions of the network. Although many additional centrality measures exist, we focused on these three metrics because they are widely used and biologically interpretable.

In the next step, network nodes were converted into a Boolean representation indicating the presence or absence of each gene (included in the networks) within each sample. Based on this binary representation, the similarity between samples was evaluated using the Jaccard similarity metric. Specifically, the Jaccard distance was computed to obtain a pairwise distance matrix, where each entry represents the dissimilarity between two samples. The distance matrix was then transformed into a similarity matrix by computing *S* = 1 − *D*, ensuring that higher values correspond to greater similarity between samples. The Jaccard similarity metric was chosen because it provides a natural measure of overlap between gene sets while remaining robust to differences in network size across samples.

A *k*-nearest neighbor graph was then constructed by selecting, for each sample, the *k* = 5 most similar samples. The corresponding similarity values were inserted into an adjacency matrix, thereby creating weighted edges between each node and its nearest neighbors. The adjacency matrix was subsequently symmetrized to ensure that the resulting graph was undirected. Community detection on this *k*-nearest neighbor graph was performed using both the Louvain algorithm [14] and the Leiden algorithm [15].

For the Louvain algorithm, implemented using the python-louvain module [16], the resolution parameter was varied between 0.5 and 1.5 and seed values ranging from 0 to 3000 were tested. The partition with the highest modularity was selected as the optimal partition. Repeating the procedure across multiple seeds allowed us to verify that the resulting community structure was stable. The resulting partitions were also visually inspected to confirm that connections within communities were stronger than those between communities. Additionally, we examined the gene composition of each community to verify that the clusters exhibited distinct molecular profiles.

To perform Leiden community detection, the NetworkX graph was converted into an igraph object [17] while preserving node names and edge weights. As in the Louvain analysis, seed values between 0 and 3000 were explored and the partition with the highest modularity score was selected.

Next, we determined the proteins associated with each cluster based on the samples assigned to that cluster. These proteins were filtered using Fisher’s exact test, retaining only those with a false discovery rate (FDR) below 0.05. Proteins were subsequently selected as potential markers if their Youden index exceeded the mean Youden index plus one standard deviation.

To characterize each cluster, enrichment analysis was performed on the corresponding gene sets. For this purpose, we used the “GO_Biological_Process_2025” library from EnrichR [18]. For each cluster, the resulting list of enriched terms was filtered to retain only unique entries with an FDR lower than 0.05. We then examined the presence or absence of the previously identified marker genes within the identified biological processes.

To further characterize the detected communities, we constructed a consensus network for each community by integrating the interactions observed across the samples belonging to that community. In these networks, edge weights represented the number of cases in which a given interaction appeared relative to the total number of cases. For each of the six resulting networks, we identified influential nodes by calculating degree, betweenness centrality, and eigenvector centrality, allowing us to detect potential hubs within each community. Because the edge weights reflected interaction frequency, they were converted into distances using the transformation *distance* = 1/*weight* in order to perform centrality calculations that rely on path lengths.

Finally, we computed a composite score to estimate the overall importance of nodes within each network. Interestingly, proteins previously identified as specific to certain biological processes did not appear among the highest-ranked nodes. This outcome may be advantageous, as targeting extremely central nodes could lead to increased toxicity. Therefore, hubs were defined as nodes with scores greater than the mean plus one standard deviation. The remaining proteins were subsequently queried in DrugBank [19] to determine whether existing drugs target them. Even when no known drugs were identified, these proteins may still represent promising candidates for further investigation.

## 3 Results

We constructed the networks using only proteins encoded by genes that were either mutated or differentially expressed. The number of nodes per network ranges from 563 to 3,851, with most networks containing approximately 2,700–3,500 nodes. The number of interactions varies between 1,243 and 22,114, with the majority of networks containing approximately 7,000–18,000 interactions. The large number of differentially expressed genes observed in this analysis is consistent with the extensive transcriptional dysregulation reported in glioblastoma. Although applying a more stringent threshold could have reduced the number of detected genes, we intentionally retained the current cutoff in order to capture a broader set of proteins within the resulting interaction networks. This choice ensures that potentially relevant nodes are represented in the networks, even if some exhibit moderate levels of differential expression. Including such nodes at this stage was considered preferable to introducing, in later analytical steps, proteins encoded by genes that do not show differential expression. However, not all proteins were captured in the networks: approximately 60% of down-regulated proteins, 50% of up-regulated proteins, and 75% of proteins encoded by mutated genes were included. The networks are summarized in Figure 2.

**Figure 2:**
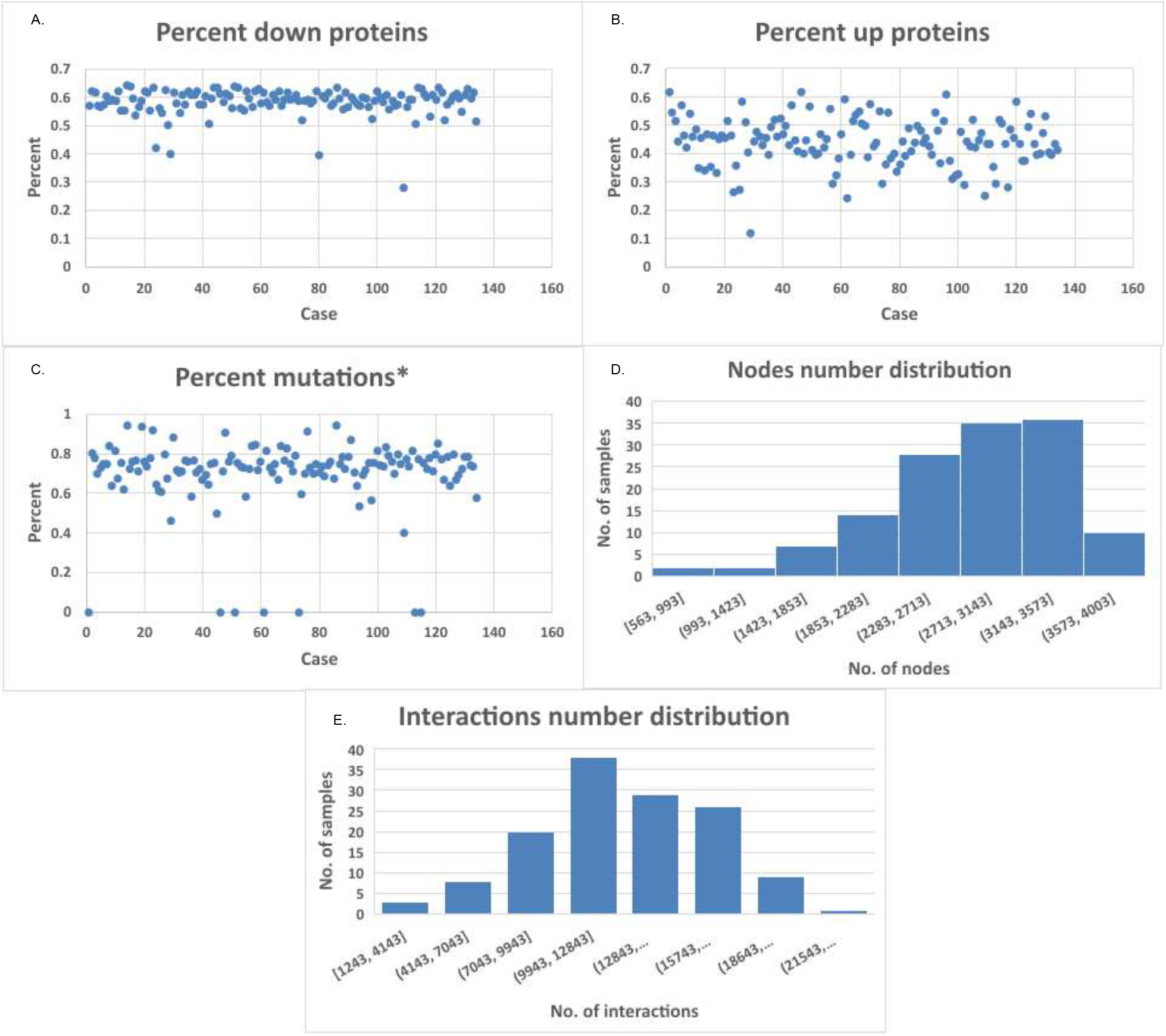
Network statistics. **A.** The percent of down-regulated proteins caught in each network.**B**. Percent of up-regulated proteins caught in each network. **C**. Percent of proteins encoded by genes that express mutations caught in each network; *samples lacking initial mutations were assigned the value 0.**D**. The distribution of the number of network based on the number of nodes. **E**. The distribution of the number of network based on the number of interactions.

Across these networks, we identified the top 10 hubs that were consistently ranked highly across the three centrality measures (degree, betweenness centrality, and maximal clique centrality): ETS1, PIK3R1, TP53, AR, YBX1, REST, MYC, EGFR, DISC1, and YWHAZ. All identified genes were queried in DisGeNET [20], and each has previously been reported to be associated with glioblastoma.

Notably, the hub most frequently identified across the different centrality measures appears among the top five nodes for each measure in only approximately half of the cases.

To investigate potential groupings of tumors, we constructed a network representing similarities among all samples. The resulting network contains 134 nodes and 570 edges. We applied the Louvain community detection algorithm and selected the partition with the highest modularity score (0.5165), indicating a strong community structure in the sample similarity network. The same modularity score was obtained when applying the Leiden algorithm. Both algorithms produced nearly identical community assignments, differing only in the ordering of the communities and the placement of a single sample. To determine the most appropriate assignment for this sample, we calculated the sum of the edge weights connecting it to neighboring nodes in each community. The results indicated that the sample was more strongly connected to the community identified by the Leiden algorithm. The Leiden optimal partition comprised six communities containing 30, 27, 26, 21, 15, and 15 samples, respectively. The communities identified by the Leiden algorithm are illustrated in Figure 3. The visualization for the Louvain algorithm would be nearly identical.

**Figure 3:**
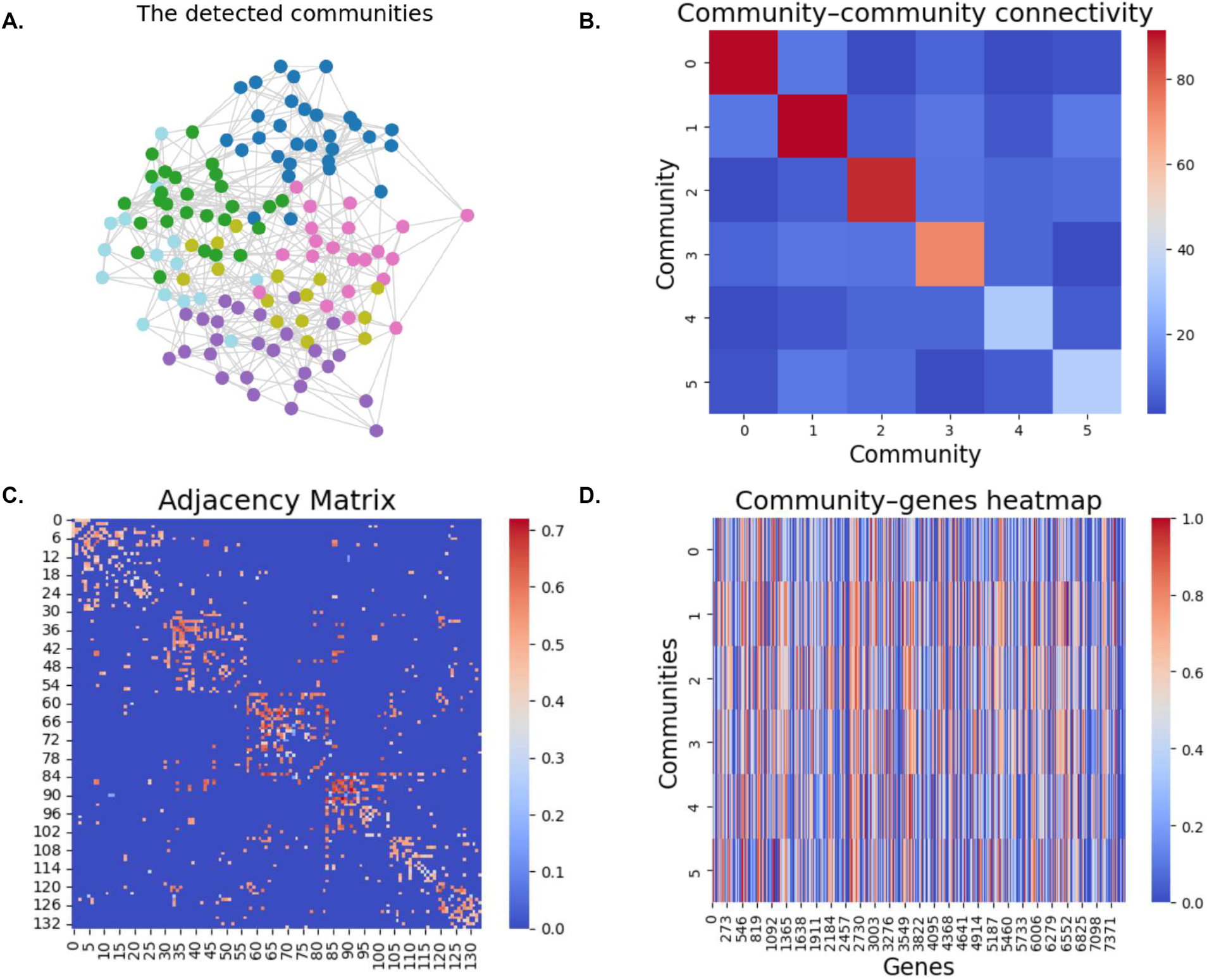
The detected communities based on Leiden algorithm. **A.** The network of samples with each community colored differently. **B**. The connectivity structure across the six communities captures the relative density of intra-community connections versus inter-community connections.**C**. The adjacency matrix for the 134 samples, capturing the pairwise connectivity structure between samples. **D**. The gene expression intensity for each gene across the six inferred clusters.

Since the two algorithms produced nearly identical results, and the only discrepant sample was ultimately assigned according to the Leiden algorithm, all subsequent analyses follow the community ordering obtained from the Leiden partition. After applying Fisher’s exact test with an FDR threshold of 0.05, the numbers of genes associated with each community were: 350 for cluster 0, 465 for cluster 1, 1072 for cluster 2, 283 for cluster 3, 27 for cluster 4, and 4 for cluster 5. Because some genes were shared across multiple clusters (specifically 126 duplicates), we further investigated which genes could be considered cluster-specific markers. Table 1 reports the number of genes specific to each cluster and lists the five highest-ranked genes according to the Youden index.

**Table 1:**
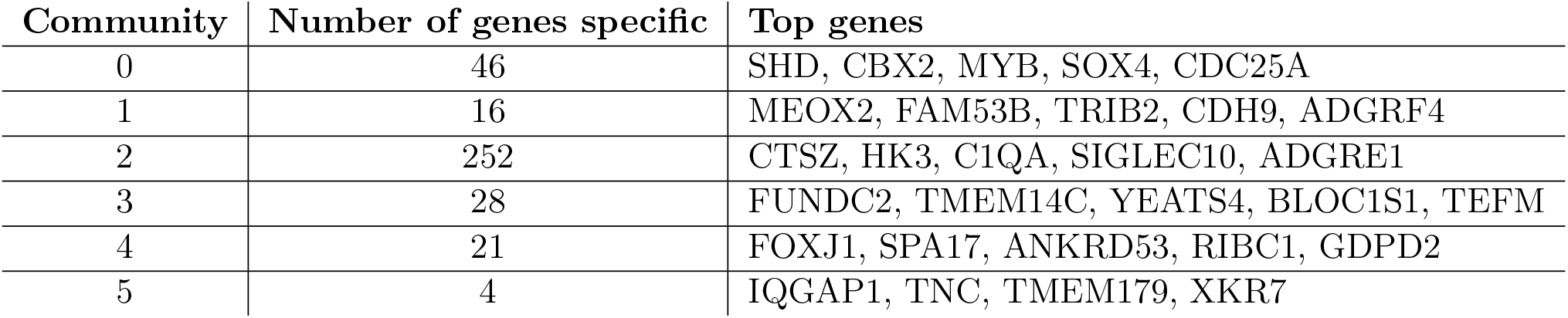
Genes specific for each community.

| Community | Number of genes specific | Top genes |
| --- | --- | --- |
| 0 | 46 | SHD, CBX2, MYB, SOX4, CDC25A |
| 1 | 16 | MEOX2, FAM53B, TRIB2, CDH9, ADGRF4 |
| 2 | 252 | CTSZ, HK3, C1QA, SIGLEC10, ADGRE1 |
| 3 | 28 | FUNDC2, TMEM14C, YEATS4, BLOC1S1, TEFM |
| 4 | 21 | FOXJ1, SPA17, ANKRD53, RIBC1, GDPD2 |
| 5 | 4 | IQGAP1, TNC, TMEM179, XKR7 |

None of the top five identified markers correspond to the top hubs previously identified at the dataset level. This is expected, as hubs—although not present in every sample—tend to occur across a larger number of samples than the cluster-specific markers.

For each community, enrichment analysis was performed on the genes identified through Fisher’s exact test. The analysis revealed multiple biological processes associated with each community. The three most significant processes for each community, ranked by false discovery rate (FDR), are presented in Table 2. These results suggest that the six communities correspond to distinct functional profiles.

**Table 2:** The number of previously identified potential marker genes that are included in the resulting biological processes, along with the top three biological processes identified.

| Community | No. of genes involved / no. of genes in the community | Biological processes |
| --- | --- | --- |
| 0 | 28/350 | DNA Damage Response (GO:0006974)<br>Regulation of G2/M Transition of Mitotic Cell Cycle (GO:0010389)<br>Cytoskeleton-Dependent Cytokinesis (GO:0061640) |
| 1 | 5/465 | Negative Regulation of Fibroblast Growth Factor Receptor Signaling Pathway (GO:0040037)<br>Mitotic DNA Replication (GO:1902969)<br>Regulation of Fibroblast Growth Factor Receptor Signaling Pathway (GO:0040036) |
| 2 | 179/1072 | Inflammatory Response (GO:0006954)<br>Positive Regulation of Cytokine Production (GO:0001819)<br>Cytokine-Mediated Signaling Pathway (GO:0019221) |
| 3 | 3/283 | Cytoplasmic Translation (GO:0002181)<br>Macromolecule Biosynthetic Process (GO:0009059)<br>Gene Expression (GO:0010467) |
| 4 | 12/27 | Axonemal Dynein Complex Assembly (GO:0070286)<br>Axoneme Assembly (GO:0035082)<br>Determination of Left/Right Symmetry (GO:0007368) |
| 5 | 2/4 | Engulfment of Apoptotic Cell (GO:0043652)<br>Glomerular Epithelial Cell Development (GO:0072310)<br>Podocyte Differentiation (GO:0072112) |

Community 0 is enriched for processes related to genomic maintenance and neuronal cellular functions, suggesting potential alterations in mechanisms responsible for preserving genomic stability. Community 1 is associated with pathways involved in cell proliferation and growth regulation, including processes linked to fibroblast growth factor receptor (FGFR) signaling. Community 2 shows enrichment in inflammatory and cytokine-mediated signaling pathways, which are consistent with known roles of immune and microglial activity in the glioblastoma tumor microenvironment. Community 3 is enriched for processes involved in macromolecule biosynthesis, gene expression, and cytoplasmic translation, reflecting the increased biosynthetic activity required to sustain rapid tumor growth. Community 4 is associated with processes related to cytoskeletal organization and axoneme assembly, suggesting potential roles in cellular motility and structural organization. Finally, community 5 includes processes related to epithelial cell differentiation and barrier-associated mechanisms, which may be relevant for interactions with the blood–brain barrier and tissue homeostasis.

Taken together, these results indicate that the detected communities capture distinct biological aspects of glioblastoma progression. These aspects are differentially intensified across communities and include proliferative signaling, inflammatory responses, biosynthetic activity, cellular organization, and genomic maintenance.

Given that these biological characteristics differ across communities, we next investigated the central hubs within each community in order to identify potential candidate targets. Hubs were identified using a composite score derived from three centrality measures: degree, betweenness centrality, and eigenvector centrality. These measures capture complementary aspects of node importance, including local connectivity, control over information flow, and influence within the network structure.

Interestingly, the highest-ranked hubs did not coincide with the top candidate markers, and conversely, the top markers were not necessarily among the most central nodes in the networks.

Nevertheless, within each community we identified genes that were simultaneously recognized as markers and occupied central positions in the corresponding networks. These genes may represent particularly interesting candidates, as they combine cluster specificity with structural importance in the interaction networks. The results are summarized in Table 3; for community 2 we report only the first 15 positions out of a total of 40.

**Table 3:** Genes that are hubs in each community and, at the same time, involved in biological processes and potential markers.

| Community | Genes |
| --- | --- |
| 0 | TFAP2A, MAD2L1, TOP2A, MYB, ESPL1, SGO1, CDC25A, CLSPN, MCM10, PKMYT1 and MYBL2 |
| 1 | PTN, EPHB4 |
| 2 | SPI1, CEBPB, BCL3, SYK, SHC1, BATF, TGFBR2, IL16, TGFBR1, HCK, TNFAIP3, ITGB2, TLR2, ANXA2, CD53 |
| 3 | SNAPIN, LAMTOR5 |
| 4 | ENKD1 |
| 5 | IQGAP1 |

Some of the identified genes correspond to known drug targets according to DrugBank [19]. For example, TGFBR1 (community 2) is targeted by galunisertib, an experimental drug currently under clinical investigation for glioblastoma. Other drugs are primarily used or studied in different cancer types. For instance, bosutinib, used in leukemia, targets HCK (community 2), while dasatinib, also used in leukemia, acts on EPHB4 (community 1). In addition, cerdulatinib, investigated for T-cell lymphoma, targets SYK (community 2). Interestingly, the antimalarial drug artenimol also interacts with targets identified in our analysis, including IQGAP1 (community 5) and ANXA2 (community 2).

Several genes identified as important in distinguishing communities currently lack known drug targets, including ENKD1, SNAPIN, and LAMTOR5. According to the Human Protein Atlas [21], SNAPIN is expressed in several cancer types, while LAMTOR5 shows moderate expression in multiple cancers and is highly expressed in the choroid plexus. ENKD1 exhibits weak to moderate expression in cancers and is strongly expressed in the flocculonodular lobe of the cerebellum.

## 4 Discussion / Future work

In this study, we investigated whether network-based analysis can help stratify glioblastoma tumors into biologically meaningful groups. By integrating differential expression and mutation data with protein–protein interaction information, we constructed patient-specific molecular networks and derived a similarity network across tumors. Community detection applied to this network revealed six tumor communities characterized by distinct gene compositions and biological processes. These results suggest that network-based representations of molecular alterations can capture aspects of glioblastoma heterogeneity that may not be apparent when analyzing individual genes in isolation.

Our analysis reinforces the value of integrating network topology with functional interpretation, a perspective well established in the literature, with seminal contributions including [22]. While highly central hubs tend to represent broadly connected proteins present across many tumors, cluster-specific markers capture more localized molecular signatures. The genes that simultaneously display central network roles and cluster specificity may therefore represent particularly interesting candidates for further biological investigation. Several of the genes identified in this study correspond to known drug targets, supporting the biological relevance of the detected communities and suggesting potential opportunities for therapeutic exploration.

Despite these encouraging results, we note several limitations.

First, the analysis relies on protein–protein interaction data derived from a global human interactome, which may not fully capture context-specific interactions present in glioblastoma cells. In future work, we will address this limitation by incorporating context-specific interaction data, in particular datasets derived from glioblastoma or relevant neural cell types as they become available. We will also investigate filtering strategies based on GBM-specific gene expression and complementary multi-omics data to retain biologically relevant interactions. These improvements are expected to enhance the biological relevance of the reconstructed networks and increase the robustness and interpretability of downstream analyses.

Second, the patient similarity network is constructed using a binary representation of gene presence, which simplifies quantitative expression information. Alternative similarity measures that incorporate expression magnitude or pathway activity may further refine the resulting clusters. In fact, we explored an alternative approach by constructing a sample network directly from gene expression values. Specifically, we restricted the analysis to genes identified through differential expression. Genes with mutations were not included unless they were also differentially expressed, as those not meeting this criterion were assumed to exhibit relatively uniform expression levels across samples and thus provide limited discriminatory power. The modularity values and corresponding number of clusters obtained across all 3,000 seeds used in the Leiden algorithm are presented in the Figure 4.

**Figure 4:**
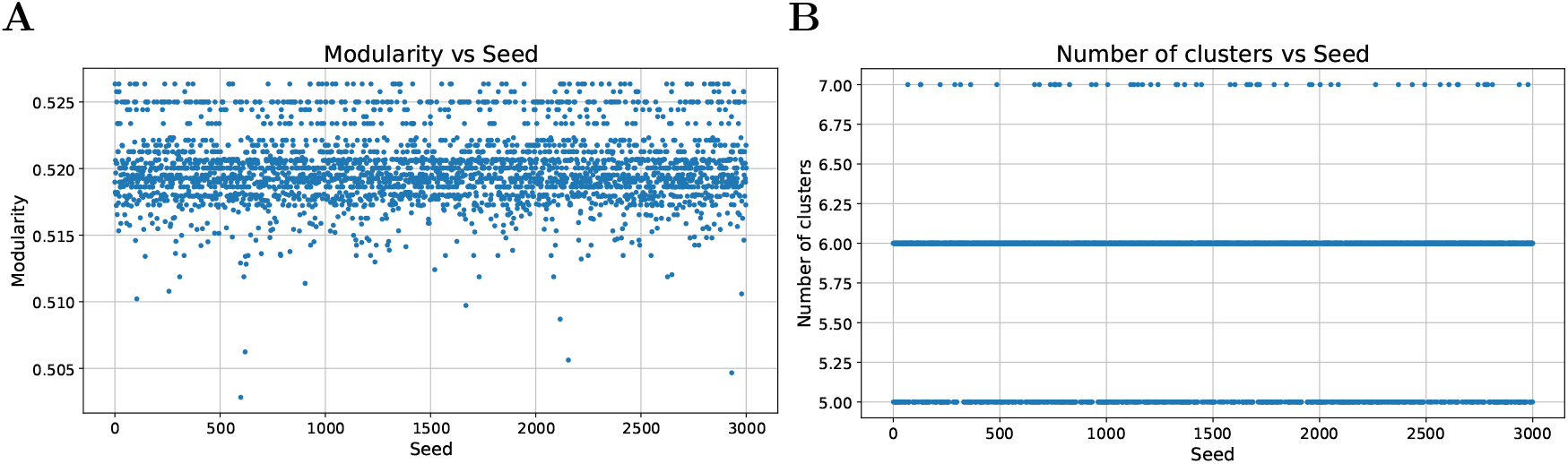
The modularity and the number of clusters when taking into account the expression values.**A.** The modularity obtained for the 3000 seeds. **B**. The number of clusters for the same 3000 seeds.

Third, our analysis relies on a single dataset. We explored additional sources and identified [23], which provides two separate mRNA sequencing datasets: one for control samples and another for mutation data. However, it was not possible to integrate mutation profiles with differentially expressed genes at the level of individual cases. Consequently, we analyzed only the differentially expressed genes, treating each dataset independently as provided. In one dataset, 140 cases of primary glioblastoma were identified within a cohort of 693 samples, whereas the second dataset contained only 85 such cases out of 325 samples. We then constructed sample networks using the same methodology and analyzed them with the Leiden algorithm. The results revealed six communities in the larger dataset and five in the smaller one (Figure 5). It remains unclear whether the lower number of communities is due to the smaller cohort size or the absence of mutation data in the analysis.

**Figure 5:**
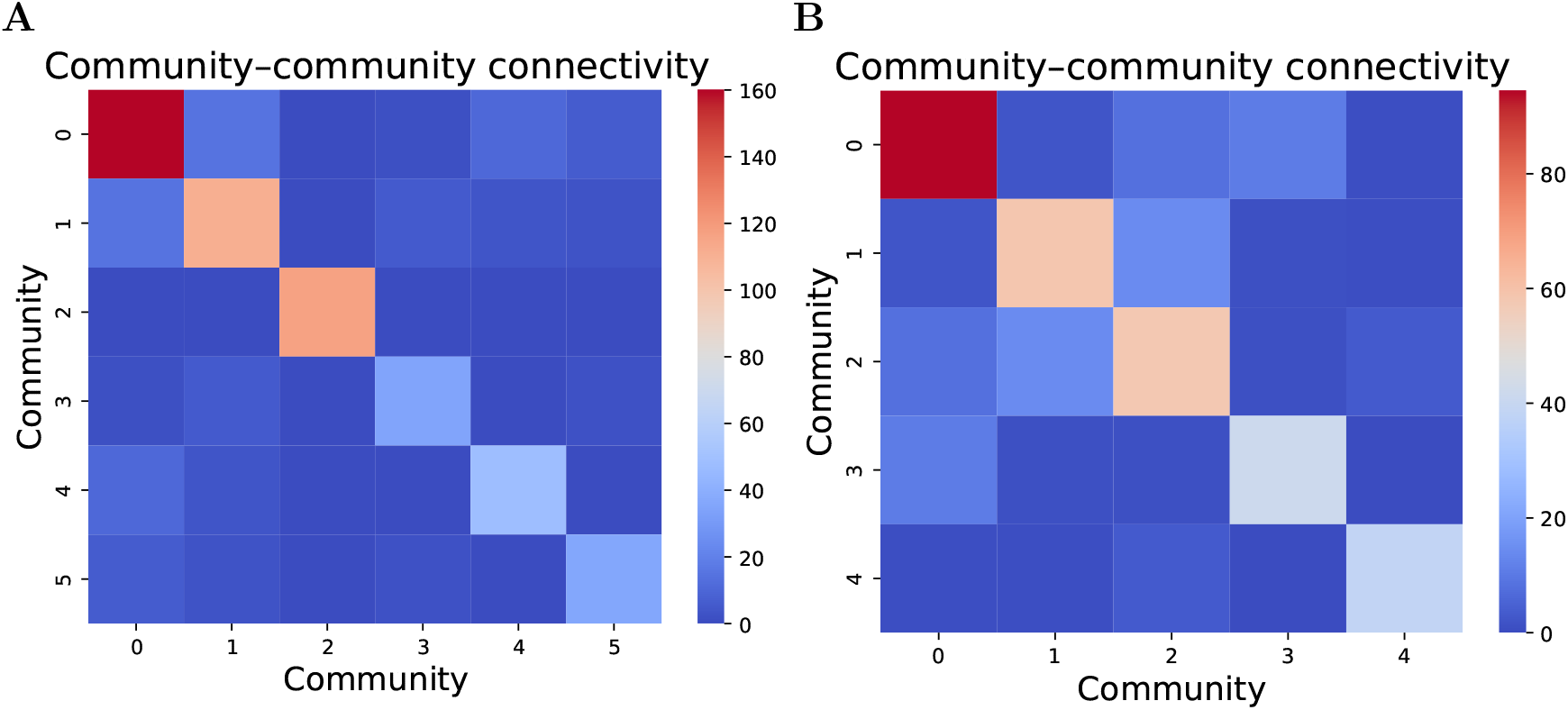
Connectivity between communities. Communities are shown on each axis and we can see that the number of those inside the community is greater than that of those outside it. **A.** In the network made on 693 dataset. **B**. In the network made on 325 dataset.

Fourth, the method relies on several fine-tuned parameters and threshold choices, which introduce approximations and can lead to differences in the results. For example, when constructing the sample network, we selected k=5 nearest neighbors. However, modularity tends to decrease as k increases, since the network becomes denser. From this perspective, the highest modularity would be achieved at k=2. In contrast, if we focus on instability, a larger value such as k=7 may be preferable (see Figure 6). Examining the number of communities further supports this trade-off: starting from k=5, the rate of change (slope) decreases gradually, suggesting a more stable community structure beyond this point.

**Figure 6:**
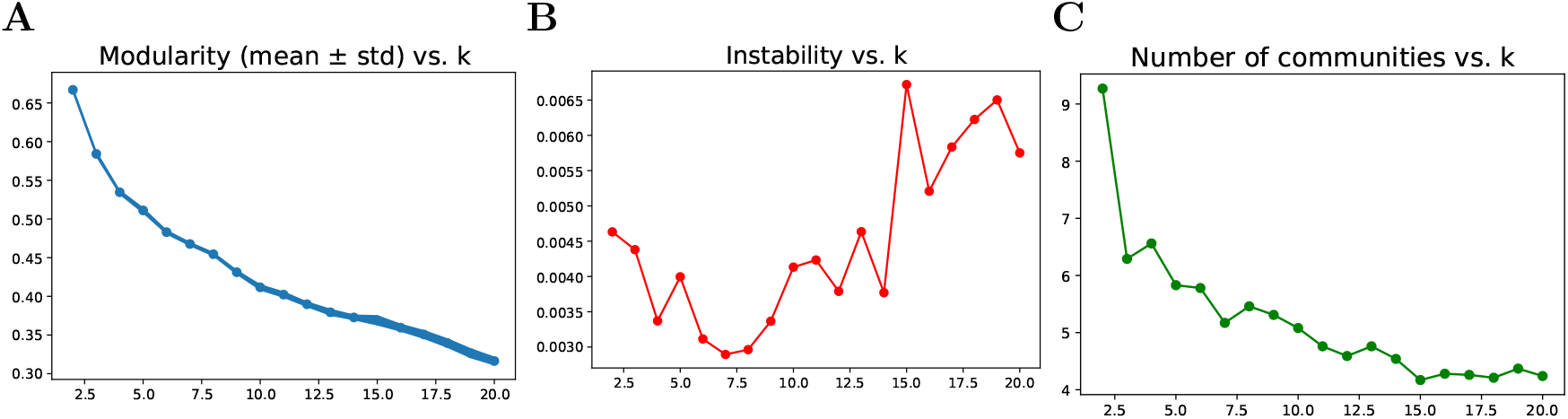
Variation of modularity, instability, number of communities depending on the number of neighbors. **A.** Modularity. **B**. Instability. C Number of communities.

Fifth, we agree that comparing the identified communities with the established glioblastoma subtypes reported by [3]. would provide valuable biological validation of our results. However, such a comparison was not feasible within the current study due to the lack of consistently available subtype annotations for the datasets used, as well as differences in data preprocessing and feature selection.

Moreover, our analysis was designed as an unsupervised, network-based exploration aimed at identifying intrinsic community structure without imposing prior subtype labels. As such, the six communities identified here should be interpreted as data-driven groupings that may partially overlap with known subtypes but are not constrained by them. Mapping these communities to established molecular subtypes represents an important direction for future work, which could further clarify the biological relevance and clinical interpretability of the detected structures. The biological interpretation of the detected communities is currently based solely on computational inference and it requires experimental validation to confirm its biological validity.

Future work could extend this framework in several directions. Incorporating additional molecular layers, such as epigenetic data, copy-number variation, or single-cell transcriptomics, may provide a more comprehensive representation of tumor heterogeneity. In addition, integrating clinical variables such as treatment response and survival outcomes could help assess the potential clinical relevance of the identified communities. Ultimately, combining network-based stratification with functional and clinical data may contribute to a more detailed understanding of glioblastoma biology and support the development of more personalized therapeutic strategies.

## 5 Code availability

Our code is available at https://github.com/nicoletasiminea/GBM_communities.

## 6 Acknowledgments

This study was supported by the Ministry of Research, Innovation and Digitalization through the National Recovery and Resilience Plan (PNRR) of Romania, Pillar III, Component C9/Investment no. 8 (I8) - PNRR–III-C9–2023–I8, contract no 760096, ID proiect – CF 68/23.05.2023.

This work was performed through the Core Program within the National Research, Development and Innovation Plan 2022-2027, carried out with the support of MRID, project no. 2302101 (SIA-PRO), contract no 7N/2022.

This work was carried out partially within the “Large Language Models for the European Union (LLMs4EU)”, project no. 101198470, call DIGITAL-2024-AI-B-06-LANGUAGE, funded by the European Union. Views and opinions expressed are however those of the author(s) only and do not necessarily reflect those of the European Union or the European Commission. Neither the European Union nor the granting authority can be held responsible for them.

## 7 Contributions

N.S. contributed to the original idea, method development, design process, data collection, data processing, analysis of results, biological interpretation of results, and manuscript preparation. D.F. contributed to the biological interpretation of results and manuscript preparation. M.P. contributed to analysis of results, manuscript preparation, and funding acquisition. A.P. contributed to the original idea, design process, analysis of results, manuscript preparation, and funding acquisition. I.P. contributed to the original idea, method development, design process, analysis of results, manuscript preparation, and funding acquisition.

## References

[1] Pouyan, A. et al. Glioblastoma multiforme: insights into pathogenesis, key signaling pathways, and therapeutic strategies. Molecular Cancer 24, 58 (2025). URL 10.1186/s12943-025-02267-0.

[2] Weller, M. et al. European Association for Neuro-Oncology (EANO) guideline on the diagnosis and treatment of adult astrocytic and oligodendroglial gliomas. The Lancet Oncology 18, e315–e329 (2017). URL 10.1016/S1470-2045(17)30194-8.

[3] Verhaak, R. G. et al. Integrated Genomic Analysis Identifies Clinically Relevant Subtypes of Glioblastoma Characterized by Abnormalities in PDGFRA, IDH1, EGFR, and NF1. Cancer Cell 17, 98–110 (2010). URL 10.1016/j.ccr.2009.12.020.

[4] Tanvir, R. B. & Mondal, A. M. Cancer Biomarker Discovery from Gene Co-expression Networks Using Community Detection Methods. In 2019 IEEE International Conference on Bioinformatics and Biomedicine (BIBM), 2097–2104 (2019).

[5] Sun, Q. et al. Identification of candidate biomarkers for GBM based on WGCNA. Scientific Reports 14, 10692 (2024). URL 10.1038/s41598-024-61515-3.

[6] Zhou, J. et al. Construction of co-expression modules related to survival by WGCNA and identification of potential prognostic biomarkers in glioblastoma. Journal of Cellular and Molecular Medicine 25, 1633–1644 (2021). URL 10.1111/jcmm.16264.

[7] Foltyn-Dumitru, M. et al. Cluster-based prognostication in glioblastoma: Unveiling heterogeneity based on diffusion and perfusion similarities. Neuro-Oncology 26, 1099–1108 (2023). URL 10.1093/neuonc/noad259.

[8] Zhang, G. et al. A Novel Molecular Classification Method for Glioblastoma Based on Tumor Cell Differentiation Trajectories. Stem Cells International 2023, 2826815 (2023). URL 10.1155/2023/2826815.

[9] Tomczak, K., Czerwińska, P. & Wiznerowicz, M. Review The Cancer Genome Atlas (TCGA): an immeasurable source of knowledge. Contemporary Oncology/Współczesna Onkologia 68–77 (2015).

[10] Gysi, D. M. et al. Network medicine framework for identifying drug-repurposing opportunities for COVID-19. Proceedings of the National Academy of Sciences 118, e2025581118 (2021). URL 10.1073/pnas.2025581118.

[11] Robinson, M. D., McCarthy, D. J. & Smyth, G. K. edgeR: a Bioconductor package for differential expression analysis of digital gene expression data. Bioinformatics 26, 139–140 (2009). URL 10.1093/bioinformatics/btp616.

[12] Durinck, S. et al. BioMart and Bioconductor: a powerful link between biological databases and microarray data analysis. Bioinformatics 21, 3439–3440 (2005). URL 10.1093/bioinformatics/bti525.

[13] Hagberg, A. A., Schult, D. A. & Swart, P. J. Exploring Network Structure, Dynamics, and Function using NetworkX. Python in Science Conference (2008). URL 10.25080/TCWV9851.

[14] Blondel, V. D., Guillaume, J.-L., Lambiotte, R. & Lefebvre, E. Fast unfolding of communities in large networks. Journal of Statistical Mechanics: Theory and Experiment 2008, P10008 (2008). URL 10.1088/1742-5468/2008/10/P10008.

[15] Traag, V. A., Waltman, L. & van Eck, N. J. From louvain to leiden: guaranteeing well-connected communities. Scientific Reports 9 (2019). URL 10.1038/s41598-019-41695-z.

[16] Aynaud, T. python-louvain x.y: Louvain algorithm for community detection. https://github.com/taynaud/python-louvain (2020).

[17] Csardi, G. & Nepusz, T. The igraph software. Complex syst 1695, 1–9 (2006).

[18] Kuleshov, M. V. et al. Enrichr: a comprehensive gene set enrichment analysis web server 2016 update. Nucleic Acids Research 44, W90–W97 (2016). URL 10.1093/nar/gkw377.

[19] Knox, C. et al. DrugBank 6.0: the DrugBank Knowledgebase for 2024. Nucleic Acids Research 52, D1265–D1275 (2023). URL 10.1093/nar/gkad976.

[20] Piñero, J. et al. DisGeNET: a discovery platform for the dynamical exploration of human diseases and their genes. Database 2015, bav028 (2015). URL 10.1093/database/bav028.

[21] Pontén, F., Jirström, K. & Uhlen, M. The Human Protein Atlas-a tool for pathology. The Journal of Pathology 216, 387–393 (2008). URL 10.1002/path.2440.

[22] Jeong, H., Mason, S. P., Barabási, A.-L. & Oltvai, Z. N. Lethality and centrality in protein networks. Nature 411, 41–42 (2001). URL 10.1038/35075138.

[23] Zhao, Z. et al. Chinese Glioma Genome Atlas (CGGA): A Comprehensive Resource with Functional Genomic Data from Chinese Glioma Patients. Genomics, Proteomics & Bioinformatics 19, 1–12 (2021). URL https://www.sciencedirect.com/science/article/pii/S1672022921000450.

